# Glutaric aciduria type 3 is a naturally occurring biochemical trait in inbred mice of 129 substrains

**DOI:** 10.1101/2020.11.11.378463

**Authors:** João Leandro, Aaron Bender, Tetyana Dodatko, Carmen Argmann, Chunli Yu, Sander M. Houten

## Abstract

The glutaric acidurias are a group of inborn errors of metabolism with different etiologies. Glutaric aciduria type 3 (GA3) is a biochemical phenotype with uncertain clinical relevance caused by a deficiency of succinyl-CoA:glutarate-CoA transferase (SUGCT). SUGCT catalyzes the succinyl-CoA-dependent conversion of glutaric acid into glutaryl-CoA preventing urinary loss of the organic acid. Here, we describe the presence of a GA3 trait in mice of 129 substrains due to SUGCT deficiency, which was identified by screening of urine organic acid profiles obtained from different inbred mouse strains including 129S2/SvPasCrl. Molecular and biochemical analyses in an F2 population of the parental C57BL/6J and 129S2/SvPasCrl strains (B6129F2) confirmed that the GA3 trait occurred in *Sugct*^129/129^ animals. We evaluated the impact of SUGCT deficiency on metabolite accumulation in the glutaric aciduria type 1 (GA1) mouse model. We found that GA1 mice with SUGCT deficiency have decreased excretion of urine 3-hydroxyglutaric acid and decreased levels glutarylcarnitine in urine, plasma and kidney. Our work demonstrates that SUGCT contributes to the production of glutaryl-CoA under conditions of low and pathologically high glutaric acid levels. Our work also highlights the notion that unexpected biochemical phenotypes can occur in widely used inbred animal lines.

**Take home message:** Glutaric aciduria type 3 is a naturally occurring trait in mice of the 129 substrains

## Introduction

Glutaric acid is a dicarboylic acid derived from glutaryl-CoA [1]. Glutaric aciduria is a biochemical condition characterized by increased excretion of glutaric acid in the urine. Degradation of lysine, but also hydroxylysine and tryptophan are the main sources of glutaryl-CoA and therefore also of glutaric acid. Three distinct types have been described. Glutaric aciduria type 1 (GA1; MIM #231670) is a dangerous neurometabolic disease caused by a deficiency in glutaryl-CoA dehydrogenase (GCDH). Glutaric aciduria type 2 (GA2; MIM #231680) is also known as multiple acyl-CoA dehydrogenase deficiency and can present as a severe multisystem disorder. GA2 is caused by a defect in the alpha or beta subunit of the electron transfer flavoprotein, or the electron transfer flavoprotein dehydrogenase. These proteins are involved in oxidizing the FADH_2_ cofactor from dehydrogenases by shuttling electrons to ubiquinone, and play a role in fatty acid, amino acid and choline metabolism. Glutaric aciduria type 3 (GA3; MIM 231690) is caused by a deficiency of succinyl-CoA:glutarate-CoA transferase (SUGCT).

In contrast to GA1 and GA2, GA3 is considered a biochemical phenotype of questionable clinical significance [2-5]. This means that this condition can be diagnosed through biochemical and genetic methods, but is considered not harmful. Although GA3 is often diagnosed in individuals with clinical symptoms, this is likely due to ascertainment bias. GA3 has been described in up to 18 individuals, many are asymptomatic, whereas others have symptoms such as gastrointestinal problems and cyclic vomiting [2, 3, 5-8]. A causal relationship between disease and the biochemical diagnosis of GA3 has been doubted based on several observations. Most importantly, healthy cases have been identified in urine-based newborn screening as well as studies of families investigated for a symptomatic case of GA3 or an unrelated diagnosis [2, 3, 5, 7]. Furthermore, the clinical symptoms associated with GA3 are non-specific, further increasing the possibility that the biochemical abnormalities were associated by chance. This implies that the clinical symptoms in these patients are caused by other (genetic) factors. Indeed, in some cases alternative genetic causes were identified that may better explain the phenotype including β-thalassemia, 6q terminal deletion syndrome, autoimmune hyperthyroidism and an autosomal recessive *TMIE* mutation [3, 5, 6]. Some of the GA3 cases were followed over time (more than 15 years) and remained asymptomatic [2, 3, 7]. Of note, the mutation that causes GA3 in the Amish (rs137852860 [2]) has an allele frequency of 9.3% in the Amish and 0.8% in non-Finnish Europeans and the gnomAD database contains 10 individuals homozygous for this variant [9]. The presence of homozygotes in this population-based database is generally considered good evidence that a variant is not disease-causing. Therefore the current consensus in the field is that SUGCT deficiency does not lead to a clinically significant disease [4].

*SUGCT*, the gene encoding SUGCT, was identified using genetic mapping in Amish GA3 families [2]. The metabolite accumulation in GA3 patients and a *Sugct* KO mouse model suggests that SUGCT mainly catalyzes the succinyl-CoA-dependent conversion of glutaric acid into glutaryl-CoA [10, 11] (Fig. 1). Humans and mice with defective SUGCT have increased plasma and urine levels of glutaric acid. Importantly and in contrast to GA1 patients, they do not have elevated 3-hydroxyglutaric acid and glutarylcarnitine (C5DC). Urinary glutaric acid excretion further increases in GA3 patients after lysine loading clearly establishing a role in endogenous lysine metabolism [3, 5]. Although SUGCT is not part of the canonical linear lysine degradation pathway, these observations suggest that a significant fraction of glutaryl-CoA undergoes spontaneous [12] or enzyme-mediated hydrolysis to glutaric acid before further metabolism by GCDH can take place (Fig. 1). The action of SUGCT prevents carbon loss due to urinary glutaric acid excretion and can be viewed as metabolite repair [13]. SUGCT is a mitochondrial enzyme, which is not consistent with the initial suggestion that GA3 was caused by a defect in a presumed peroxisomal glutaryl-CoA oxidase [3, 5]. The mitochondrial localization of SUGCT, however, has been unequivocally established by biochemical characterization of a mitochondrial dicarboxyl-CoA:dicarboxylic acid CoA transferase (EC 2.8.3.13), the presence of a mitochondrial targeting sequence in the SUGCT protein, the mitochondrial localization of a SUGCT-GFP fusion protein and the inclusion in MitoCarta, which is an inventory of mammalian mitochondrial proteins [2, 11, 14-16]. A peroxisomal targeting signal could not be identified.

**Figure 1.**
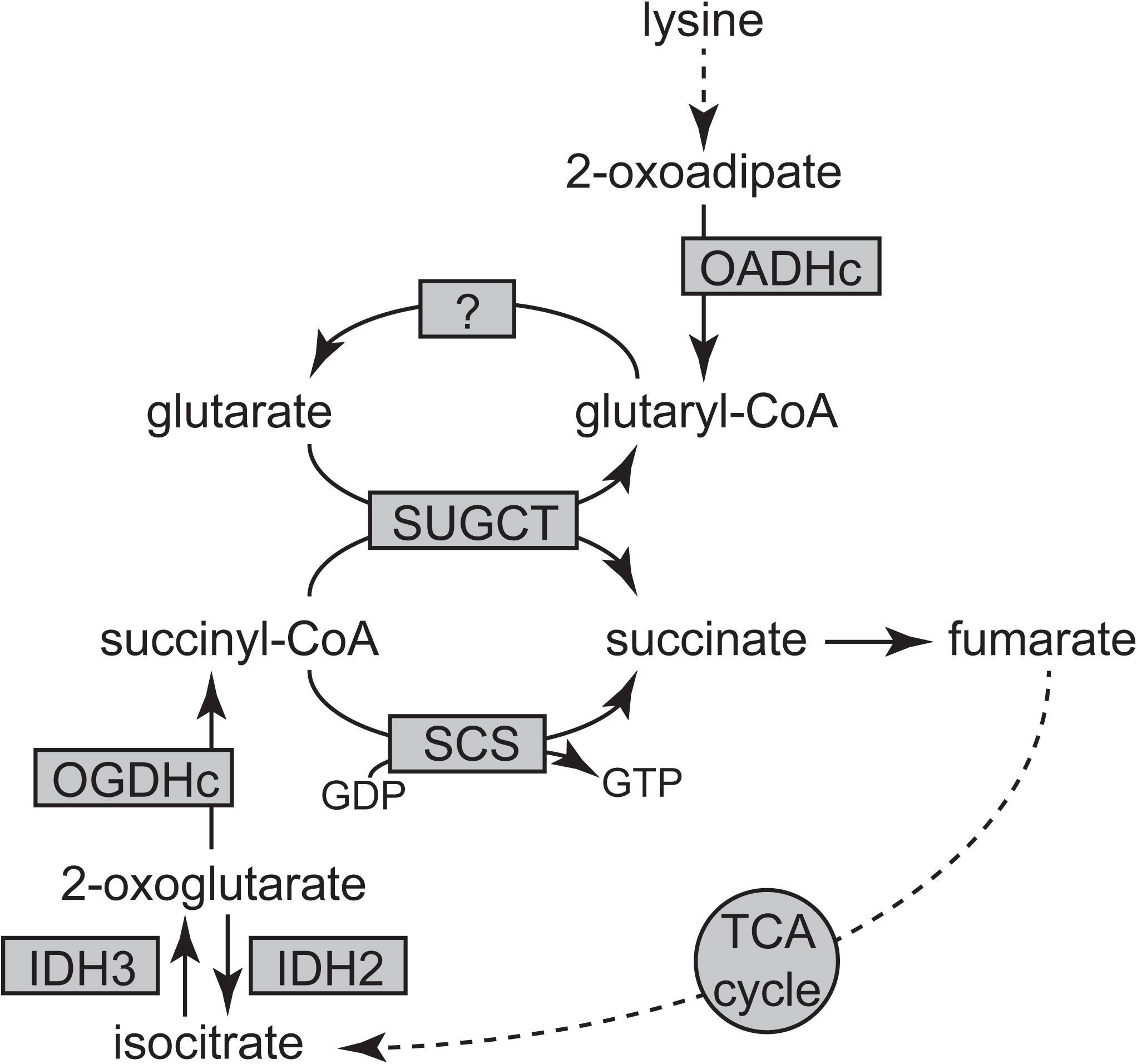
Schematic representation of glutaryl-CoA metabolism and the role of SUGCT. The interactions between lysine metabolism and the TCA cycle are displayed. The abbreviations are: IDH, isocitrate dehydrogenase; OADHc, 2-oxoadipate dehydrogenase complex; OGDHc, 2-oxoglutarate dehydrogenase complex; SCS, succinyl-CoA synthetase; SUGCT, succinyl-CoA:glutarate-CoA transferase; TCA, tricarboxylic acid.

Here we demonstrate that mice of the 129 substrain have SUGCT deficiency and display a GA3 trait. We also show that SUGCT deficiency can modulate metabolite accumulation in GA1 mice.

## Material and Methods

### Generation of the mouse models

All animal experiments were approved by the IACUC of the Icahn School of Medicine at Mount Sinai (#2016-0490) and comply with the National Institutes of Health guide for the care and use of laboratory animals (NIH Publications No. 8023, revised 1978).

Six week old male 129S2/SvPasCrl, DBA/2J and C57BL/6J mice were ordered and used for random urine sampling. B6129F1 mice were generated using male C57BL/6J and female 129S2/SvPasCrl parents. The B6129F2 cohort was generated after intercrossing of B6129F1 mice. All B6129F2 mice were genotyped for the *Sugct, D2hgdh, Dhtkd1* and *Ivd* loci.

The *Gcdh* KO model [17] is C57BL/6NJ congenic. B6129F1 mice were crossed with *Gcdh*^+/-^ mice in order to generate B6129F2 mice (with a 75% C57BL/6 background). These B6129F2 mice were used to generate mice for the experimental cohort. For this, we intercrossed the *Gcdh*^+/-^ *Sugct*^129/B6^ *Dhtkd1*^129/B6^ mice. The genotype distributions from the resulting litters are available in Table S1. The experimental cohort consisted of *Gcdh*^-/-^ mice that were grouped according to their genotype at the *Sugct* and *Dhtkd1* loci; 129/129, 129/B6 and B6/B6.

All mice were fed a standard rodent chow (PicoLab Rodent Diet 20). Urine of individual mice was collected on multiple days and pooled in order to obtain sufficient sample volume. Mice were euthanized in a random fed state (afternoon) at ∼10 weeks of age using CO_2_. Blood from the inferior vena cava was collected for EDTA plasma isolation, and organs were rapidly excised and snap frozen in liquid nitrogen and subsequently stored at −80°C.

### Mouse genotyping

The *Sugct* locus was genotyped using a restriction fragment length polymorphism (RFLP) at Chr13:17,425,964T>C. For this we amplified 155 bp of intron 9 using the following forward and reverse primers; 5’-tca gga cca ttg act cac aca-3’ and 5’-tgt gtc caa tca gca aag cc-3’. The SNP creates an ApaI site and the PCR product is cleaved for *Sugct*^129^, but not for *Sugct*^B6^. The *Ivd* locus was genotyped using a RFLP at rs47004510. For this we amplified 134 bp covering intron 8, exon 9 and intron 9 using the following forward and reverse primers; 5’-cca ctc ccc act tga caa ca-3’ and 5’-acc tgg aat tgg ccg atc tt-3’. The SNP removes a StuI site and the PCR product is cleaved for *Ivd*^B6^, but not for *Ivd*^129^. Genotyping of the *D2hgdh* locus was complicated by the presence of two duplications containing the majority of the *D2hgdh* gene. Ultimately, we genotyped rs30957932, a SNP in the promoter region. For this we amplified a 307 bp fragment using the following forward and reverse primers; 5’-gct ggg gga tgc tga agt aa-3’ and 5’-gtc cat cag cag aca cga ct-3’. The SNP removes an EcoNI site and the PCR product is cleaved for *D2hgdh*^B6^, but not for *D2hgdh*^129^. The *Dhtkd1* locus was genotyped as described [18, 19].

The *Gcdh* transgenic locus was genotyped using a common forward (Exon 1 *Gcdh* 5’-gaa cca atg agc aac ccc ta-3’) and two allele-specific reverse primers (wild type: Exon 3 *Gcdh* 5’-cag ttt ctc atc ggc agt ca-3’, and mutant: LacZ 5’-gac agt atc ggc ctc agg aa-3’). The wild type allele yields a 713 bp product, whereas the *Gcdh* KO allele a product of ∼400 bp.

### Immunoblotting

Frozen mouse tissues (kidney, liver and brain) were homogenized in ice-cold RIPA buffer supplemented with protease inhibitor cocktail (∼20 mg tissue per 1 mL lysis buffer) using a TissueLyser II apparatus (Qiagen) and the homogenates were centrifuged at 1,000xg, 10 min at 4°C. Total protein was determined using the BCA method. Proteins were separated on a Bolt(tm) 4–12% Bis-Tris Plus Gel, blotted onto a nitrocellulose membrane and detected using the following primary antibodies: SUGCT (NBP2-69820, Novus Biologicals) and citrate synthase [GT1761] (GTX628143, Genetex) or [N2C3] (GTX110624; RRID:AB_1950045, Genetex). Proteins were visualized using IRDye 800CW or IRDye 680RD secondary antibodies (LI-COR, 926-32210; RRID:AB_621842, 926-68070; RRID:AB_10956588, 926-32211; RRID:AB_621843, and 926-68071; RRID:AB_10956166) in an Odyssey CLx Imager (LI-COR) with Image Studio Lite software (version 5.2, LI-COR). Equal loading was checked by Ponceau S staining and the citrate synthase signal. We also tested another SUGCT (NBP1-98273, Novus Biologicals) antibody, but this one was not reactive with mouse SUGCT in immunoblotting.

### RNA sequencing and RT-PCR

Total RNA from kidney was isolated from 4 wild type and 4 *Ehhadh* KO mice (congenic C57BL/6) for an unrelated research project. At the same time, RNA was also isolated from 1 DBA/2J and 1 129S2/SvPasCrl kidney sample. RNA was submitted to the Genomics Core Facility at the Icahn Institute and Department of Genetics and Genomic Sciences for further processing. mRNA-focused cDNA libraries were generated using Illumina reagents (polyA capture), and samples were run on an Illumina HiSeq 2500 sequencer to yield a read depth of approximately 56 million 100 nucleotide single end reads per sample. Reads from fastq files were aligned to the mouse genome mm10 (GRCm38.75) with STAR (release 2.4.0g1 [20]) and summarized to gene- and exon-level counts using featureCounts [21]. *Sugct* was differentially expressed between wild type and *Ehhadh* KO mice, but the change in expression was small (logFC=-0.281, adj P value = 0.046). We therefore reused all these data to study differences in exon level expression of *Sugct* between the 129S2/SvPasCrl sample and all other samples.

cDNA was synthesized from total RNA isolated from liver of 129S2/SvPasCrl, DBA/2J and C57BL/6J mice. A *Sugct* cDNA fragment was amplified using the following forward (exon 1: 5’-gct ctc gct ggg gtt gag-3’) and reverse (exon 6: 5’-aca cat cac aaa tgg ctg caa-3’) primers. A *Rplp0* cDNA fragment was amplified as control (5’-atg ggt aca agc gcg tcc tg-3’ and 5’-gcc ttg acc ttt tca gta ag-3’).

### Metabolite analyses

Plasma acylcarnitines, urine acylcarnitines and urine organic acids were measured by the Mount Sinai Biochemical Genetic Testing Lab (now Sema4). Urine creatinine was measured by Jaffe’s reaction. Urine organic acids were quantified using a standard curve and pentadecanoic acid as internal standard. Tissue acylcarnitines were measured in freeze‐dried samples after derivatization to propylesters followed by analysis on an Agilent 6460 Triple Quadrupole Mass Spectrometer [22, 23]. All animal and metabolite data are available in Table S2 and S3.

### Statistical analyses

Data are displayed as the mean ± the standard deviation (SD) as indicated in the figure legends. Differences between groups of mice were evaluated using one-way analysis of variance (ANOVA) with Turkey’s multiple comparison test, a Kruskal-Wallis test or a two-way ANOVA as indicated. Significance is indicated in the figures. Allele effect sizes were determined using linear regression analysis. All analyses were performed in GraphPad Prism 6.

## Results

### Glutaric aciduria type 3 in mice of the 129 substrain

We analyzed the urine organic acid profile in randomly collected urine samples from C57BL/6J, 129S2/SvPasCrl and DBA/2J mice. Previously, we described the elevated 2-oxoadipic acid levels in urine of C57BL/6J mice [19]. Here, we report increased 2-hydroxyglutaric acid in C57BL/6J and increased glutaric acid in 129S2/SvPasCrl (Fig. 2A). Levels of 3-hydroxyglutaric acid were undetectable in all samples. Although 129S2/SvPasCrl mice are known to have increased C5-carnitine [19], no other abnormalities in the acylcarnitine profile were noted in this strain. Specifically, C5DC levels were normal.

**Figure 2.**
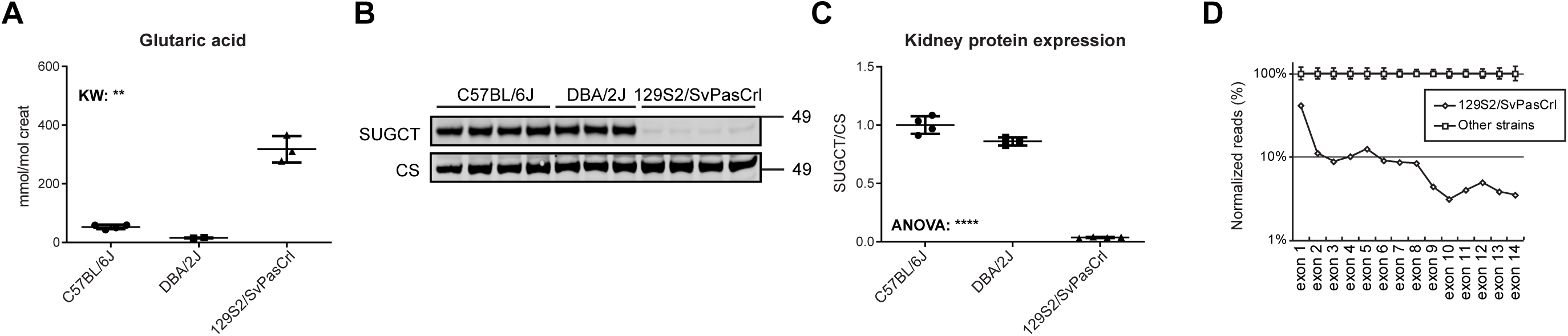
Identification of a GA3 trait and SUGCT deficiency in mice of the 129 substrain. (A) Quantification of glutaric acid in urine samples from C57BL/6J (n = 4 of which one pooled sample), DBA/2J (n = 2 pooled samples) and 129S2/SvPasCrl (n = 3 of which one pooled sample) mice. P value indicates the result of a Kruskal-Wallis test. **, P < 0.01. (B) Immunoblot analysis and (C) quantification of SUGCT protein levels in kidney of C57BL/6J, DBA/2J and 129S2/SvPasCrl mice. The P value indicates the result of an ANOVA test. ****, P < 0.0001. SUGCT expression was normalized using citrate synthase (CS). The average SUGCT/CS for C57BL/6J was set to 1. (D) Expression of *Sugct* exons. For each sample exonic *Sugct* reads were normalized to total mapped reads. The 129S2/SvPasCrl reads were then expressed as a percentage of the average reads in samples of the other strains. 129S2/SvPasCrl included RNAseq data from 1 kidney. The other strains included 8 C57BL/6 and 1 DBA/2J kidney samples. Error bars indicate SD.

We used quantitative proteomics data on liver of the 8 founder laboratory strains of the Collaborative Cross, which include C57BL/6J and 129S1/SvlmJ to investigate the cause of the glutaric aciduria [24]. Protein abundance data confirmed several previously described mild inborn errors of metabolism including deficiencies of DHTKD1, BCKDHB, D2HGDH, COX7A2L and NNT in C57BL/6J, and deficiencies of IVD and GLYCTK in 129S1/SvlmJ (Fig. S1A) [19]. In addition, we noted a deficiency of SUGCT in 129S1/SvlmJ mice. We subsequently confirmed SUGCT deficiency in kidney homogenates of 129S2/SvPasCrl mice (Fig. 2B). Residual SUGCT protein expression was 4% compared to C57BL/6J (Fig. 2C). A genetic mechanism underlying SUGCT deficiency in 129S1/SvlmJ mice is suggested by the identification of a *Sugct cis*-eQTL in liver [24]. In addition, mapping of liver expression data from the hybrid mouse diversity panel, which includes the 129×1/SvJ strain, reveals a *cis*-eQTL and significant association with many SNPs within the chromosomal location of *Sugct* (e.g. rs29516766; chr13:17,604,658; P = 3.8291e-11, Fig. S1B) [25, 26].

In order to investigate the molecular mechanism of the SUGCT defect in 129 mice, we compared RNAseq data from kidneys of mice with different genetic backgrounds. We found that while the expression level of exon 1 was fairly normal in 129S2/SvPasCrl (41%), reads mapping to downstream exons decreased first to 10% and then to 4% of control values (Fig. 2D). RT-PCR on liver cDNA revealed decreased *Sugct* expression in 129S2/SvPasCrl when compared C57BL/6J and DBA/2J (Fig. S1C). Combined these data suggest that 129 mice carry a GA3 trait.

### Association of the GA3 trait with the Sugct^129/129^ genotype in a B6129F2 population

In order to unequivocally establish that 129 mice carry a GA3 trait, we generated an F2 population of the parental C57BL/6J and 129S2/SvPasCrl strains (B6129F2) and genotyped the *Sugct* as well as the *Ivd* and *D2hgdh* loci. We measured plasma acylcarnitine and urinary organic acids in this B6129F2 cohort (Table S2). Consistent with previously reported associations [19, 27], plasma C5-carnitine was elevated in *Ivd*^129/129^ animals and urine 2-hydroglutaric acid was elevated in *D2hgdh*^B6/B6^ animals (Fig. S2A). *Sugct*^129/129^ animals showed a pronounced glutaric acid accumulation in urine, whereas levels were low in *Sugct*^B6/129^ and *Sugct*^B6/B6^ animals (Fig. 3A). Similar to GA3 in humans, there was no detection of urine 3-hydroxyglutaric acid and normal plasma C5DC (Fig. 3B). Consistent with publicly available *Sugct* mRNA expression data, immunoblotting of kidney, liver and brain homogenates showed high expression of SUGCT in kidney, lower levels in liver, and virtually undetectable levels in brain. Protein levels of SUGCT in kidney, liver and brain were deficient in *Sugct*^129/129^ animals and reduced in *Sugct*^B6/129^ animals when compared to *Sugct*^B6/B6^ animals (Fig. 3C). Next, we used linear regression analysis to estimate the effect size of the *Sucgt*^129^ allele on SUGCT protein expression in the kidney. We found that for each *Sucgt*^129^ allele, SUGCT protein levels decrease by ∼48% (Fig. 3D). SUGCT protein levels in *Sucgt*^129/129^ kidneys was 3% when compared *Sucgt*^B6/B6^ kidneys (Fig. S2B). The characterization of B6129F2 mice demonstrates that mice of the 129 substrain have an autosomal recessive GA3 trait that associates with the *Sugct*^129^ locus.

**Figure 3.**
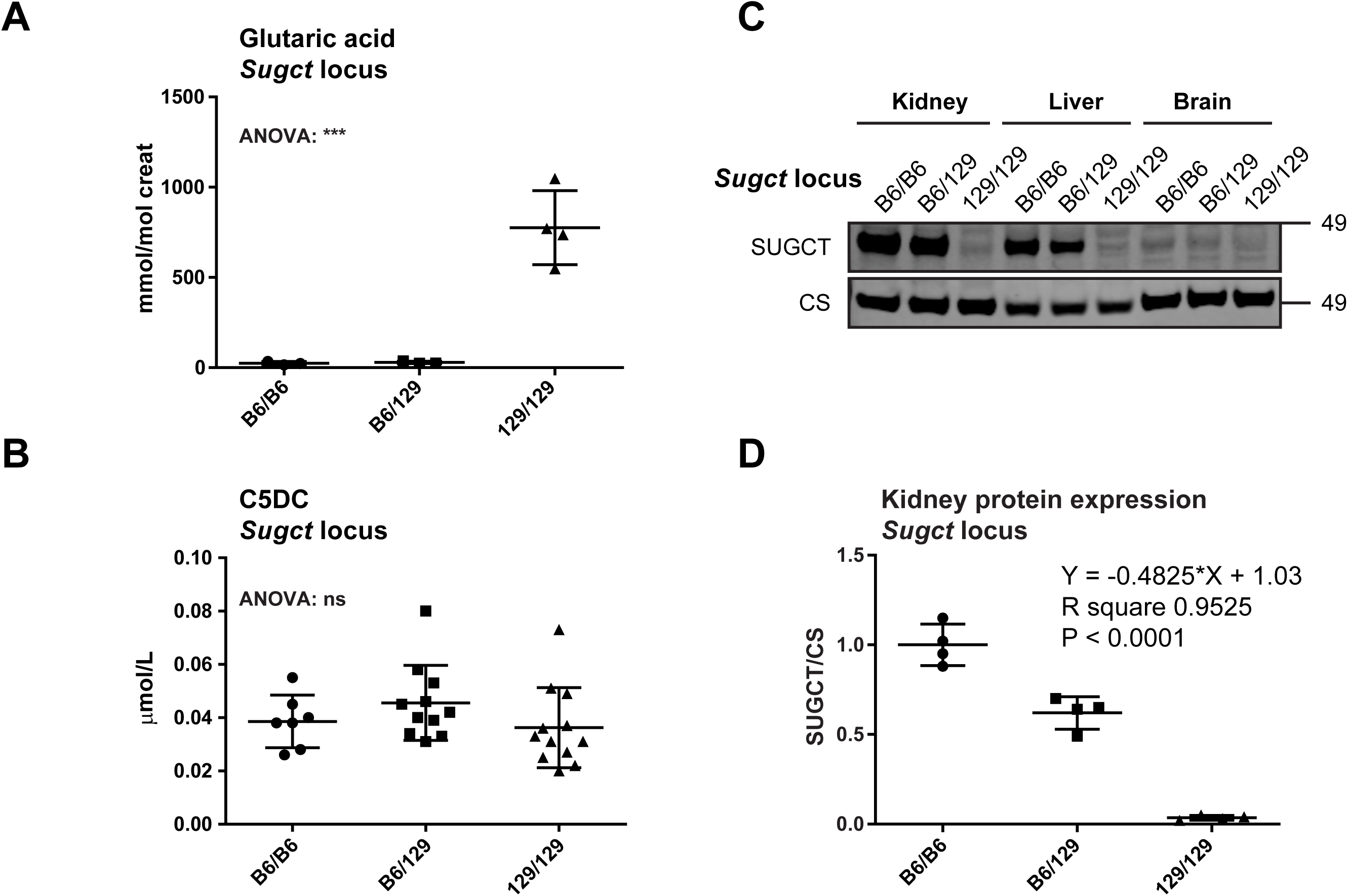
Association of the GA3 trait with the *Sugct*^129/129^ genotype in a B6129F2 population. Quantification of urine glutaric acid (A) and plasma C5DC (B) in samples of B6129F2 mice with *Sugct*^B6/B6^, *Sugct*^B6/129^ and *Sugct*^129/129^ genotypes. The P value indicates the result of an ANOVA test. ***, P < 0.001. (C) Immunoblot analysis of SUGCT in kidney, liver and brain in samples of B6129F2 mice with *Sugct*^B6/B6^, *Sugct*^B6/129^ and *Sugct*^129/129^ genotypes. Citrate synthase (CS) served as a loading control. (D) Quantification of SUGCT protein levels in kidney of B6129F2 mice with *Sugct*^B6/B6^, *Sugct*^B6/129^ and *Sugct*^129/129^ genotypes. SUGCT expression was normalized using citrate synthase (CS). The average SUGCT/CS for C57BL/6J was set to 1. Linear regression analysis was performed to estimate the effect size of the *Sugct*^129^ allele on protein expression. Error bars indicate SD.

The generation and characterization of a *Sugct* KO mouse model was recently reported [10]. Metabolomic analysis of kidney not only revealed elevated levels of glutaric acid, but also of adipic acid [10]. We evaluated levels adipic and pimelic acid in urine of the B6129F2 mice (Table S2). *Sugct*^129/129^ animals showed increased adipic acid levels in urine compared *Sugct*^B6/129^ and *Sugct*^B6/B6^ animals (Fig. S2C). Pimelic acid levels were not different between these groups.

### Sugct^129/129^ is a biochemical modifier of GA1

Genetic background is known to be a modifier of the clinical and biochemical phenotype of GA1 mice. On a mixed background (C57BL/6J x 129/SvEv), GA1 mice develop symptoms upon lysine exposure, whereas they remain asymptomatic on a C57BL/6J background [28]. These data suggest the presence of a modifier gene with the 129 allele making GA1 mice sensitive to disease, whereas the C57BL/6J allele is protective. The modifier appears to affect GA1 neuropathology through modulating substrate accumulation because the manifestation of symptoms correlates with accumulation of glutaric acid in serum, liver and brain [28, 29]. We have previously identified a functionally significant natural variant in the *Dhtkd1*^B6^ allele that causes decreased *Dhtkd1* mRNA and DHTKD1 protein, increased plasma 2-aminoadipic acid and increased urine 2-oxoadipic acid [19, 30]. This hypomorphic *Dhtkd1* allele, but also a\ *Dhtkd1* KO allele, was not protective in the GA1 mouse model [18, 31].

Next, we intercrossed the *Gcdh*^+/-^ *Sugct*^129/B6^ *Dhtkd1*^129/B6^ mice. The genotype distribution in the progeny (122 pups) was Mendelian for all combinations of *Gcdh* and *Sucgt* or *Gcdh* and *Dhtkd1* loci (Table S1). We measured plasma acylcarnitines, urine organic acids and urine acylcarnitines in all GA1 mice from this cohort (Table S3). These biochemical traits were then associated with the genotype at the *Sugct* or *Dhtkd1* loci. We found that plasma C5DC concentration was lower in *Sugct*^129/129^ animals when compared to *Sugct*^B6/129^ and *Sugct*^B6/B6^ animals (Fig. 4A). Similarly, excretion urine 3-hydroxyglutaric acid and C5DC in urine of *Sugct*^129/129^ animals was lower when compared to *Sugct*^B6/129^ and *Sugct*^B6/B6^ animals (Fig. 4A). Levels of urine glutaric acid were not different between groups (Fig. 4A). When the same mice were segregated according to their genotype at the *Dhtkd1* locus, we observed increased urine 2-oxoadipic acid in *Dhtkd1*^B6/B6^ animals, but no changes in any of the GA1 biomarkers (Fig. S3A), which confirms our previously reported work [18]. Importantly, these data demonstrate that under basal conditions, *Sugct*^129^ can act as a modifier of the biochemical phenotype of the GA1 mouse, which is in stark contrast to *Dhtkd1*^B6^.

**Figure 4.**
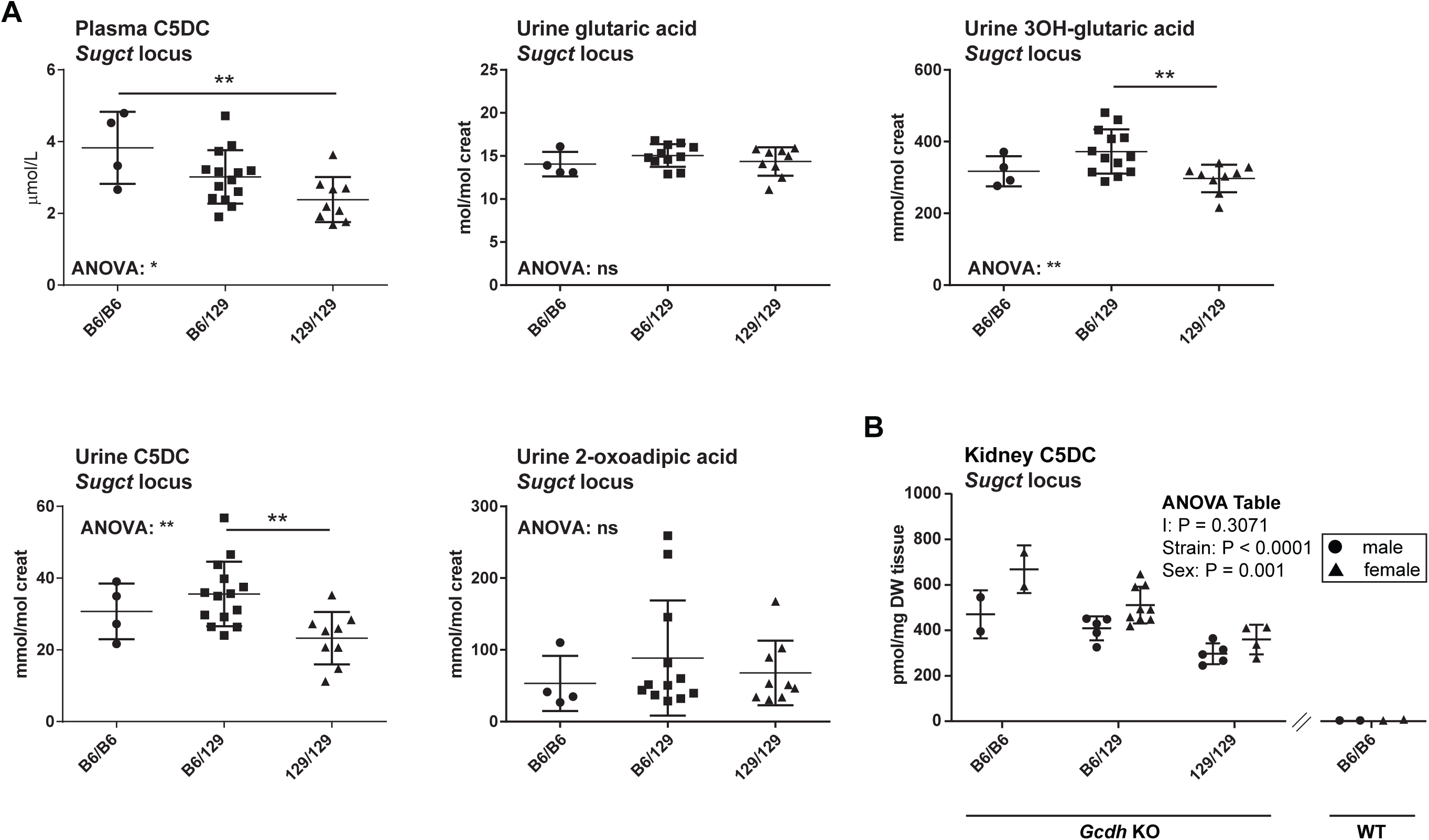
The biochemical consequences of SUGCT deficiency in GA1 mice. (A) Plasma C5DC, urine glutaric acid, 3OH-glutaric acid, C5DC and 2-oxoadipate in GA1 mice. Mice are segregated according to their genotype at the *Sugct* locus. The P values indicate the result of an ANOVA test and a Tukey’s multiple comparisons test. *, P < 0.05 and **, P < 0.01. (B) Kidney C5DC (as pmol per mg tissue dry weight (DW)) in GA1 mice. Mice are segregated according to their genotype at the *Sugct* locus. The table displays the result of a two-way ANOVA.

Urine adipic acid is elevated in GA1 mice [32]. We found that SUGCT deficiency further increased urinary excretion of adipic acid. Most of the variation, however, was explained by sex, with adipic acid levels being higher in male urine (Fig. S3B).

We measured acylcarnitines in kidney, liver and brain and analyzed levels of C5DC. In kidney, C5DC was strongly affected by *Sugct* genotype as well as sex, and displayed gene-dose dependency (Fig. 4B). C5DC was not affected by genotype in liver and brain (Fig. S3C). The tissue acylcarnitine data confirm that *Sugct*^129^ can modify the biochemical phenotype of the GA1 mouse, but the effect appears restricted to kidney, a tissue with high *Sugct* expression.

## Discussion

Here we report that mice of the 129 substrains harbor a GA3 trait caused by SUGCT deficiency. We show that the *Sugct*^129/129^ genotype causes decreased *Sugct* mRNA with ∼4% residual SUGCT protein expression leading to pronounced glutaric aciduria as well as noticeable adipic aciduria. This confirms that under physiological conditions, SUGCT mainly serves to reactivate glutaric acid into glutaryl-CoA, which prevents urinary loss and can be viewed as metabolite repair [13]. We also tested how SUGCT deficiency affects metabolite accumulation in a mouse model for GA1. We observed that GA1 mice with SUGCT deficiency have lower levels in 3-hydroxyglutaric acid and C5DC in urine and a lower concentration of plasma C5DC. In kidney, the tissue with the highest level of SUGCT expression, we observed a gene-dose dependent decrease in C5DC for the *Sugct*^129^ allele. These results indicate that even under conditions of highly elevated glutaric acid, the net flux through SUGCT proceeds towards the resynthesis of glutaryl-CoA (Fig. 1).

The increased excretion of adipic acid in *Sugct*^129/129^ mice shows that SUGCT accepts adipoyl-CoA as substrate in vivo, and indicates that there may be a mitochondrial source of adipoyl-CoA. We speculate that mitochondrial adipoyl-CoA is derived from adipoylcarnitine that may be produced by peroxisomal β-oxidation of long-chain dicarboxylic acids [33]. Adipic acid excretion is also elevated in GA1 [32, 34, 35] and urine levels increased even more with SUGCT deficiency. It is currently unknown why adipic acid excretion is increased in GA1. Potential explanations include the possibility that adipoyl-CoA is a substrate for GCDH, competitive inhibition between elevated glutaryl-CoA and adipoyl-CoA for further metabolism by other acyl-CoA dehydrogenases, or competition between glutaric acid and adipic acid for reuptake from the pre-urine in the kidney.

It is remarkable that lysine metabolism and the TCA cycle converge at multiple points around glutaryl-CoA metabolism (Fig. 1). The 2-oxoglutarate dehydrogenase complex and the 2-oxoadipate dehydrogenase complex are responsible for production of succinyl-CoA and glutaryl-CoA respectively, but have recently been demonstrated to form a hybrid complex and display overlap in substrate specificity [18, 36, 37]. Glutaric acid is a good substrate for succinyl-CoA synthetase (SCS) from the hyperthermophilic archaea *Thermococcus kodakaraensis* [38], but this has not been studied for the mammalian SCS enzymes. The reaction catalyzed by SUGCT also converts succinyl-CoA into succinate, but the CoA molecule has to be transferred onto glutaric acid (or adipic acid). Given these interactions, it would be interesting to study TCA cycle flux in the brain of the GA1 model, in particular upon high lysine feeding.

A limitation of our work is that we only studied the effect of SUGCT deficiency in GA1 mice on a regular chow diet. Under these conditions, the GA1 mouse model does not develop neurological disease. Exposure to high lysine conditions is necessary to induce a neurological phenotype. The low expression of SUGCT in the brain and the absence of significant changes in brain C5DC under basal conditions suggest that the effect of SUCGT deficiency in the brain will be limited. Given the significant pain and discomfort for the animals, we judged the high lysine studies with neurological endpoints at this stage not ethical. A second limitation is that we have not identified the causal variant that underlies GA3 in 129 mice. We found that while the expression level of exon 1 was fairly normal in kidney of 129S2/SvPasCrl, reads mapping to downstream exons decreased first to 10% and then to 4% of control values. *Sugct* is a relatively large gene (837 kbp, 0.7% of chr13) consisting of 14 exons that encode a 1,727 bp mRNA (NM_138654.3). Intron 1 (22 kbp) harbors many repetitive DNA elements due to retrotransposon insertions. Intron 1 also encodes a second poorly annotated and seemingly nonfunctional *Sugct* transcript that utilizes the same exon 1, but two alternate exons located within intron 1 (ENSMUST00000220514.1). Therefore we hypothesize that the causal variant responsible for the SUGCT defect in 129 mice, is most likely located in the intron 1 region, but the RNAseq data did not reveal any obvious aberrant splicing events. Current 129 genome sequences from the Mouse Genomes Project did not reveal any plausible causal variants, but are likely incomplete due to the low-complexity and repetitive regions in intron 1. In order to identify the variant and molecular mechanism that underlie SUGCT deficiency in the 129S2/SvPasCrl strain, future studies will need to generate a high-coverage long-read sequence dataset of a 129S2/SvPasCrl mouse, and include a characterization of all murine *Sugct* transcripts.

Our work also highlights the notion that unexpected biochemical phenotypes can occur in widely used inbred animal lines. Since these biochemical phenotypes occur in commonly used inbred strains they are not linked to deleterious phenotypes. There are at least three possible explanations for this observation. Firstly, there is often considerable residual activity of the deficient protein, which may be linked to milder or no phenotypes. This may be the case for the *Ivd* deficiency in 129 mice [19]. Secondly, some biochemical phenotypes such as GA3 are considered clinically insignificant in humans [4] and as such are also unlikely to cause disease in mice. Lastly, environmental or genetic factors, such as diet, ambient temperature or genetic modification may be necessary to induce a clinical phenotype. Given the vast number of genetically engineered mouse models that have been generated over the past decades, we speculate that genetic interactions should be expected. Indeed, we have demonstrated that a naturally occurring enzyme defect in SUGCT can modify the biochemical phenotype of a KO mouse model for GA1. Our work highlights that it is important to “mind your mouse strain” [39]. To support other investigators, we provide an updated table that summarizes naturally occurring traits in inbred mouse strains (Table S4).

## Acknowledgments

We thank Dr. Matthew Hirschey (Duke Molecular Physiology Institute, Duke University Medical Center, Durham, NC) for providing the *Gcdh* KO mice and Purvika Patel for kind and excellent technical assistance with the clinical biochemical analyses.

## Author contributions

Conception and design of the work described: SMH

Acquisition of data: JL, AB, TD, SMH

Analysis and interpretation of data: JL, CA, CY, SMH

Reporting of the work described: SMH

## Guarantor

Sander M. Houten

## Ethics statement

All animal experiments were approved by the IACUC of the Icahn School of Medicine at Mount Sinai (# IACUC-2016-0490) and comply with the National Institutes of Health guide for the care and use of Laboratory animals (NIH Publications No. 8023, revised 1978).

