## Supplementary information for "Glutaric aciduria type 3 is a naturally occurring biochemical trait in inbred mice of 129 substrains"

**Supplemental figure legends**

Figure S1. Identification of a GA3 trait and SUGCT deficiency in mice of the 129 substrain. (A) Quantitative proteomics data for liver expression of selected proteins in the 8 founder laboratory strains of the Collaborative Cross as extracted from Table S1 in Chick et al [1]. Abundance data are displayed for proteins associated with biochemical phenotypes reported in the C57BL/6J and 129S1/SvlmJ strains. This analysis confirmed several previously described mild inborn errors of metabolism including deficiencies of DHTKD1, BCKDHB, D2HGDH, COX7A2L and NNT in C57BL/6J, and deficiencies of IVD and GLYCTK in 129S1/SvlmJ. Other strains include all data from A/J, NOD/ShiLtJ, CAST/EiJ, NZO/HILtJ, WSB/EiJ and PWK/PhJ. (B) Mapping of *Sugct* (*5033411D12Rik*; Probeset ID 1421422_at) liver expression data from the hybrid mouse diversity panel using the systems genetics resource at UCLA (https://systems.genetics.ucla.edu/) [2, 3]. (C) Expression analysis of *Sugct* in liver cDNA of C57BL/6J, DBA/2J and 129S2/SvPasCrl mice. *Rplp0* served as control gene.

Figure S2. Association of the GA3 trait with the *Sugct*$^{129/129}$ genotype in a B6129F2 population. (A) Plasma C5-carnitine and urine 2-hydroglutaric acid in samples of B6129F2 mice. For plasma C5-carnitine the mice were genotyped at the *Ivd* locus, and for urine 2-hydroglutaric acid the mice were genotyped at the *D2hgdh* locus. (B) Immunoblot analysis of SUGCT in kidney samples of B6129F2 mice with *Sugct*$^{B6/B6}$, *Sugct*$^{B6/129}$ and *Sugct*$^{129/129}$ genotypes. (C) Quantification of urine adipic acid in samples of B6129F2 mice with *Sugct*$^{B6/B6}$, *Sugct*$^{B6/129}$ and *Sugct*$^{129/129}$ genotypes. The P value indicates the result of an ANOVA test. **, P< 0.01; ***, P < 0.001; ****, P< 0.0001.

Figure S3. The biochemical consequences of SUGCT deficiency in GA1 mice. (A) Plasma C5DC, urine glutaric acid, 3OH-glutaric acid, C5DC and 2-oxoadipate, and kidney C5DC in GA1 mice. Mice are segregated according to their genotype at the *Dhtkd1* locus. The P values indicate the result of an ANOVA test and a Tukey's multiple comparisons test. Ns, not significant; ****, P< 0.0001. (B) Urine adipic acid for male and female GA1 mice. Mice are segregated according to their genotype at the *Sugct* locus. (C) Liver and brain C5DC in GA1 mice. Data are displayed for the same cohort of animals (Fig. 4 and Fig. S3A), segregated according to their genotype at the *Sugct* locus. The table displays the result of a two-way ANOVA.

**Supplemental tables**

Table S1. Genotype distribution in an intercross of *Gcdh*$^{+/-}$ *Sugct*$^{129/B6}$ *Dhtkd1*$^{129/B6}$ mice. Provided as an Excel table.

Table S2. Biochemical and phenotypic data in mice of an F2 population of the parental C57BL/6J and 129S2/SvPasCrl strains (B6129F2). Provided as an Excel table.

Table S3. Biochemical and phenotypic data in the GA1 mice of an F2 population of the parental C57BL/6J and 129S2/SvPasCrl strains (B6129F2). Provided as an Excel table.

Table S4. List of selected defects due to spontaneous mutations in (commonly used) inbred mouse strains.

Table S4. List of selected defects due to spontaneous mutations in (commonly used) inbred mouse strains.

| # | Gene | Full name | Causal variant | Strain(s) | MIM | References |
|---|---|---|---|---|---|---|
| 1 | *a* | nonagouti | Many | Many | 611742 | [4] |
| 2 | *Acads* | Short-chain acyl-CoA dehydrogenase (SCAD) | Deletion Chr5:115,110,929-115,111,207 | BALB/cByJ | 201470 | [5] |
| 3 | *Alpl* | alkaline phosphatase | Unknown | C57BL/6J | 146300<br>241500<br>241510 | [6] |
| 4 | *Bckdhb* | branched chain keto acid dehydrogenase E1 subunit beta | Insertion<br>Chr9:83,942,547-83,949,878 | C57BL/6J | 248600 | [7-9] |
| 5 | *Casp4* | caspase 4, apoptosis-related cysteine peptidase | Deletion Chr9:2,328,429:2,328,433<br>skipping of exon 7<br>p.P304fsX309<br>rs235337590 | 129 substrains | | [10] |
| 6 | *Cdh23* | cadherin 23 | c.753G>A<br>in-frame skipping of exon 7 | Many strains including the C57BL/6 substrains | 601386<br>601067<br>617540 | [11] |
| 7 | *Cdt1* | chromatin licensing and DNA replication factor 1 | c.265-282delAGCCCTCCAGCTGACCCT<br>p.S89_P94del<br>rs232721151 | 129 substrains<br>LP/J | 613804 | [12] |
| 8 | *Cox7a2l* | supercomplex assembly factor I | c.216_221delGCCCAT<br>p.F72delinsLPI | C57BL/6 substrains<br>129S5/SvEvBrd<br>BALB/cJ | Not reported | [9, 13] |

| | | | | | | |
|---|---|---|---|---|---|---|
| 9 | *D2hgdh* | D-2-hydroxyglutarate dehydrogenase | Unknown | C57BL/6J | 600721 | [9]<br>This paper |
| 10 | *Dhtkd1* | dehydrogenase E1 and transketolase domain containing 1 | Insertion<br>chr2:5,924,595-5,925,150 | C57BL/6J<br>A/J<br>AKR/J<br>FVB/NJ<br>NOD/LtJ<br>NZO/HILtJ | 204750<br>245130 | [7, 8] |
| 11 | *Disc1* | disrupted in schizophrenia 1 | c.1583-1607delACCAGGCTCCCTTCCAGGTGGAGCC<br>p.Q529LfsX141 | 129 substrains<br>FVB/NJ<br>LP/J | 604906 | [14, 15] |
| 12 | *Dock2* | dedicator of cytokinesis 2 | duplication in exons 28 and 29 | C57BL/6NHsd | 616433 | [16, 17] |
| 13 | *Eci3* | enoyl-Coenzyme A delta isomerase 3 | Unknown | DBA/2J | Not applicable | [18] |
| 14 | *Glyctk* | glycerate kinase | c.196dupA<br>p.R66KfsX143<br>rs243632237 | 129 substrains<br>LP/J | 220120 | [19, 20] |
| 15 | *Gpnmb* | glycoprotein (transmembrane) nmb | c.448C>T<br>p.R150X<br>rs47598337 | DBA/2J | Not reported | [21] |
| 16 | *Ivd* | isovaleryl-CoA dehydrogenase | c.1068G>A<br>p.K356K<br>rs27440099 | 129S1/SvImJ<br>129S2/SvPasCrl<br>129P2/OlaHsd<br>129S5/SvEvBrd<br>BTBR $T^{+}$ $Itpr3^{tf}$/J<br>BUB/BnJ | 243500 | [7]<br>This paper |

| | | | | | | |
|---|---|---|---|---|---|---|
| | | | | LP/J | | |
| 17 | *Lep* | Leptin (Obese) | c.313T>C<br>p.R105X | *Ob*/*Ob* | 614962 | [22, 23] |
| 18 | *Mlycd* | malonyl-CoA decarboxylase | Insertion 5313bp Chr8:119,402,361 | DBA/2J<br>SM/J | 248360 | Our unpublished data |
| 19 | *Nnt* | nicotinamide nucleotide transhydrogenase | Deletion 17.8 kbp Chr13:119,375,448 | C57BL/6J | 614736 | [24, 25] |
| 20 | *Slc3a1* | solute carrier family 3 member 1 (rBAT) | c.1232G>A<br>p.E383K | 129S2/SvPasCrl<br>Later reported as not present in 129S1/Sv or 129S2/SvPas mice, which was confirmed by our lab | 220100 | [26] |
| 21 | *Slc11a1* | solute carrier family 11 (proton-coupled divalent metal ion transporters), member 1<br>natural resistance-associated macrophage protein 1 (*Nramp1*) | c.506A>G<br>p.D169G<br>rs47476426 | 129P2/OlaHsd<br>129S1/SvImJ<br>129S5SvEvBrd<br>C3H/HeJ<br>most other strains | | [27-31] |
| 22 | *Slc22a5* | high-affinity sodium-dependent carnitine cotransporter (OCTN2) | c.1055T>G<br>p.L352R | juvenile visceral steatosis (Jvs) | 212140 | [32] |

| | | | | | | |
|---|---|---|---|---|---|---|
| 23 | *Sugct* | succinyl-CoA:glutarate-CoA transferase | unknown | 129 substrains | 231690 | This paper |
| 24 | *Trem2* | triggering receptor expressed on myeloid cells 2 | c.TC>GA<br>p.S148E | 129S1/SvImJ<br>PWK/PhJ<br>CASA/RkJ | 618193 | |
| 25 | *Ttc7* | tetratricopeptide repeat domain 7 | An ETn early transposon insertion of 183 bp occurred in intron 14, 57 bp upstream of exon 15. | A/J | 243150 | [33] |
| 26 | *Tyr* | tyrosinase (albino / c) | c.308G>C<br>p.C103S<br>rs31191169 | A/J<br>AKR/J<br>BALB/cJ<br>FVB/NJ<br>NOD/LtJ | 203100 | [34] |

The table lists the official gene name of the affected gene, a commonly used full gene name and abbreviation, the causal variant, the strain in which the variant was first characterized, the MIM number (https://www.omim.org/) of the equivalent human genetic disorder and a selected reference. The Dec. 2011 (GRCm38/mm10) mouse assembly was used.

### Figure S1

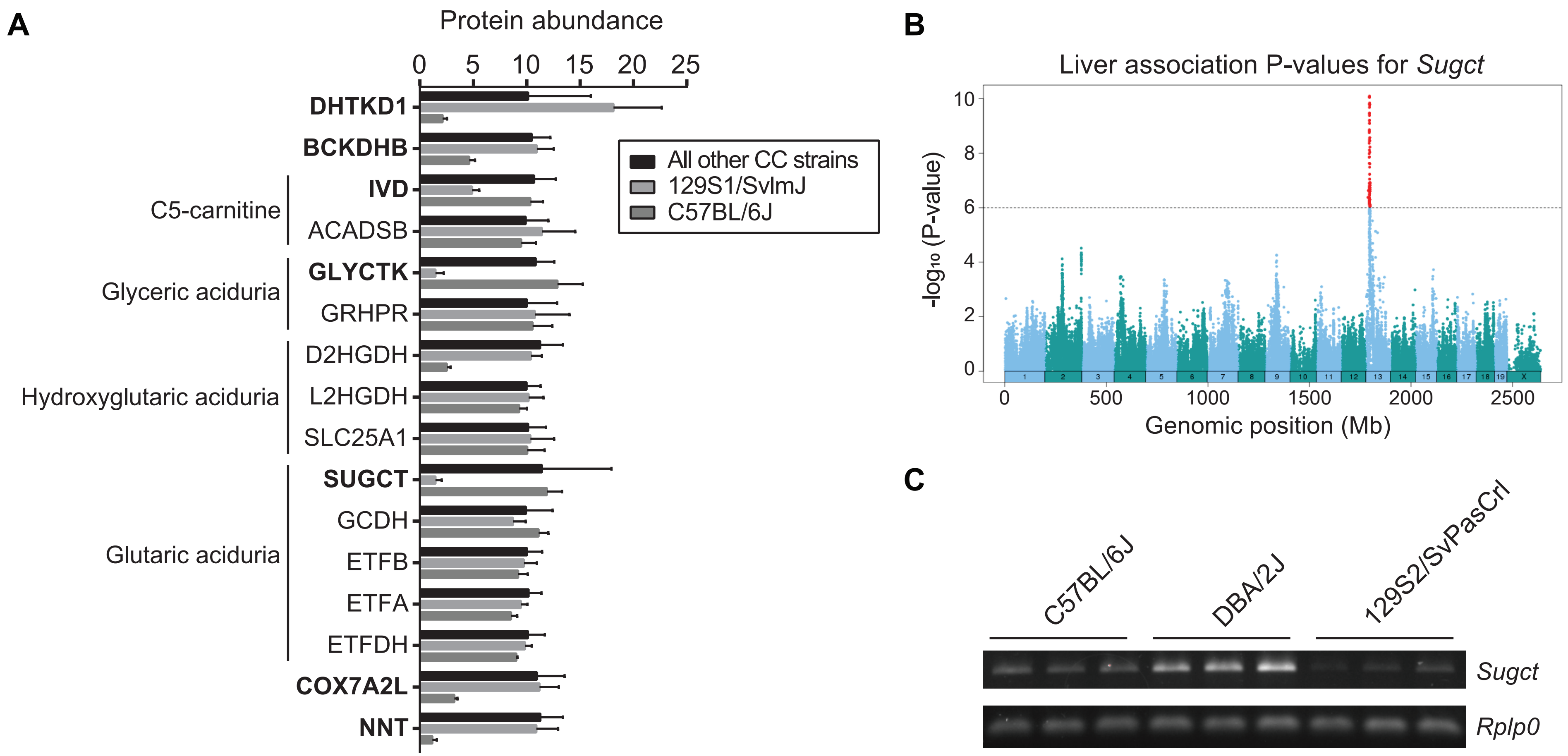
A
Protein abundance
0
5
10
15
20
25
DHTKD1
BCKDHB
IVD
ACADSB
GLYCTK
GRHPR
D2HGDH
L2HGDH
SLC25A1
SUGCT
GCDH
ETFB
ETFA
ETFDH
COX7A2L
NNT
C5-carnitine
Glyceric aciduria
Hydroxyglutaric aciduria
Glutaric aciduria
All other CC strains
129S1/SvImJ
C57BL/6J
B
Liver association P-values for Sugct
-log10 (P-value)
Genomic position (Mb)
0
500
1000
1500
2000
2500
C
C57BL/6J
DBA/2J
129S2/SvPasCrl
Sugct
Rplp0

### Figure S2

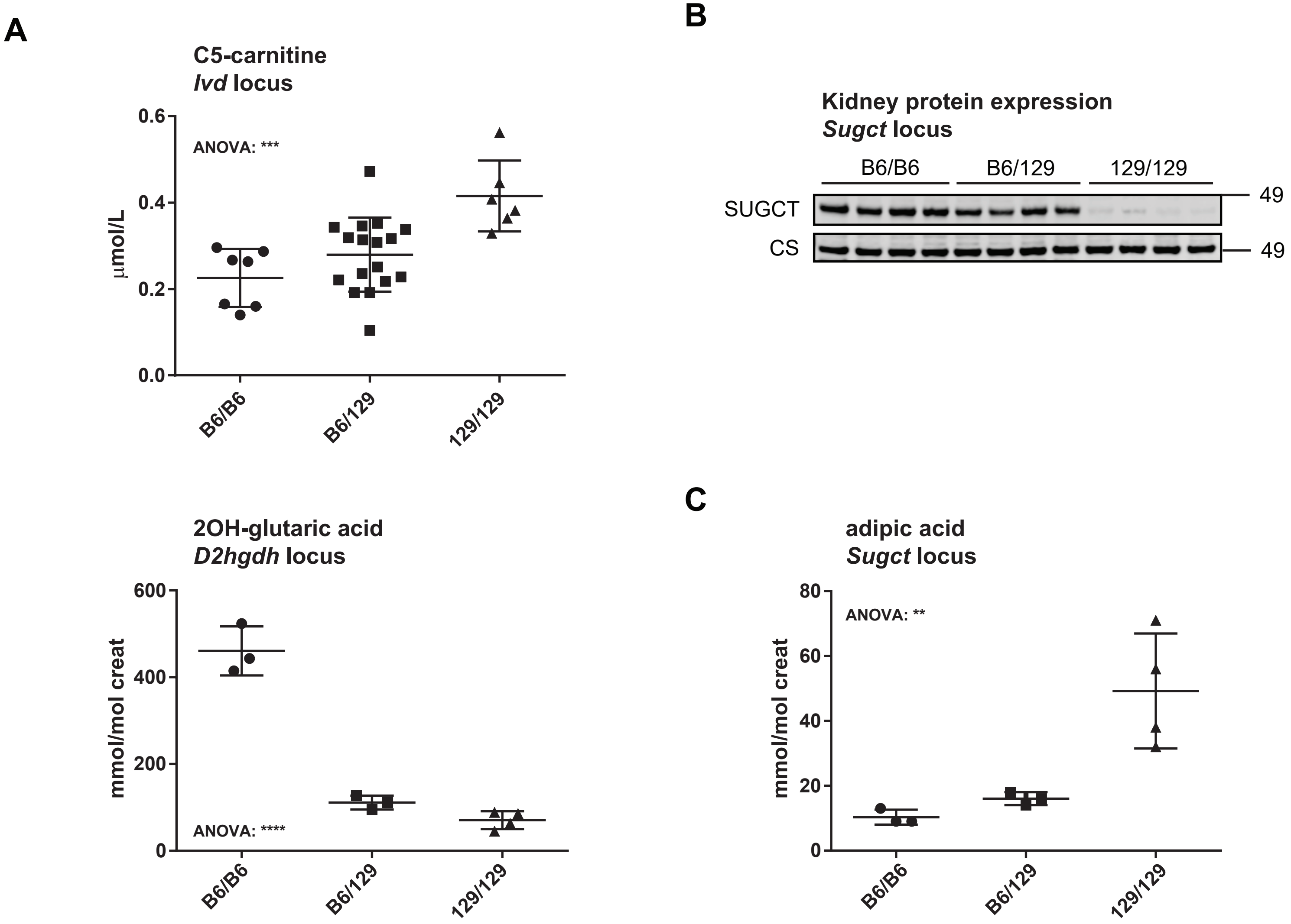
A
C5-carnitine
*Ivd* locus
ANOVA: ***
µmol/L
0.0
0.2
0.4
0.6
B6/B6
B6/129
129/129
2OH-glutaric acid
*D2hgdh* locus
ANOVA: ****
mmol/mol creat
0
200
400
600
B6/B6
B6/129
129/129
B
Kidney protein expression
*Sugct* locus
B6/B6
B6/129
129/129
SUGCT
CS
49
49
C
adipic acid
*Sugct* locus
ANOVA: **
mmol/mol creat
0
20
40
60
80
B6/B6
B6/129
129/129

### Figure S3

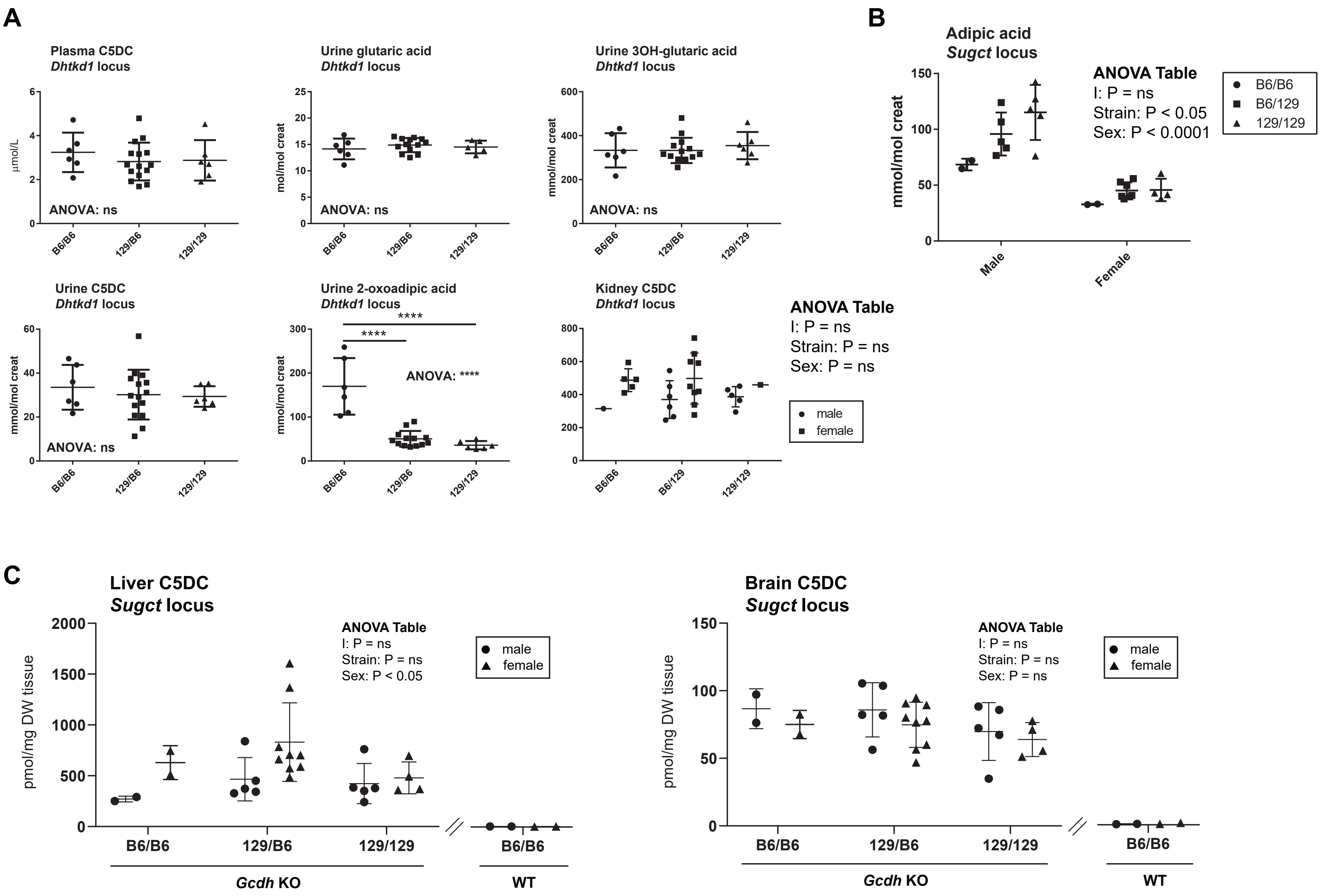
A
Plasma C5DC
Dhtkd1 locus
µmol/L
ANOVA: ns
B6/B6
129/B6
129/129
Urine glutaric acid
Dhtkd1 locus
mol/mol creat
ANOVA: ns
Urine 3OH-glutaric acid
Dhtkd1 locus
mmol/mol creat
ANOVA: ns
Urine C5DC
Dhtkd1 locus
mmol/mol creat
ANOVA: ns
Urine 2-oxoadipic acid
Dhtkd1 locus
mmol/mol creat
****
****
ANOVA: ****
Kidney C5DC
Dhtkd1 locus
B6/129
ANOVA Table
I: P = ns
Strain: P = ns
Sex: P = ns
male
female
B
Adipic acid
Sugct locus
mmol/mol creat
ANOVA Table
I: P = ns
Strain: P < 0.05
Sex: P < 0.0001
B6/B6
B6/129
129/129
Male
Female
C
Liver C5DC
Sugct locus
pmol/mg DW tissue
ANOVA Table
I: P = ns
Strain: P = ns
Sex: P < 0.05
male
female
B6/B6
129/B6
129/129
Gcdh KO
WT
Brain C5DC
Sugct locus
pmol/mg DW tissue
ANOVA Table
I: P = ns
Strain: P = ns
Sex: P = ns
male
female
Gcdh KO
WT
